# Investigating microbiota differences across chronic pancreatitis, influenced by lifestyle and genetic determinants

**DOI:** 10.1101/2025.04.04.647187

**Authors:** Abubaker Y.M Ahmed, Azita Rajai, Catherine Fullwood, Damian W Rivett, John McLaughlin, Christopher van der Gast, Ryan Marsh

## Abstract

**Background:** Chronic pancreatitis (CP) is a multifaceted and irreversible inflammatory condition that impacts both endocrine and exocrine pancreatic functions. Whilst there is evidence of a general intestinal microbial dysbiosis, differences across CP resulting from diverse triggers remains poorly elucidated.

**Methods:** In this study, we investigated how different CP triggers, namely alcohol-induced CP (AIP) and CFTR-related CP (CFRP), might undermine differences across microbiota structure and predicted function. Faecal samples from 25 CP participants were collected, alongside 9 healthy controls for microbiota analyses. Both full and short (V4) 16S rRNA sequencing was performed, including predicted functionality using PICRUSt2. Clinical metadata was integrated to investigate relationships with microbiota structure and predicted functionality with ordination-based approaches.

**Results:** Microbiota diversity patterns between groups varied slightly by sequencing approach, with differences across common core and rarer satellite taxa diversity evident. Importantly, microbiota compositional profiles were shared between approaches, with control samples clustering closer than CP samples. Between groups, significant differences in microbiota composition were apparent, with the AIP group significantly different to both control and CFRP groups across various microbiota partitions (*P <* 0.05). Notable genera drove the observed dissimilarity, including *Bifidobacterium*, *Faecalibacterium*, and *Ruminococcus*, which were decreased in relative abundance across CP. Predictive functional analyses further demonstrated increased similarity of controls as compared to CP groups. Significant variation across both microbiota structure and predicted function was explained by CP subtype (*P <* 0.01). When compared back to controls, the relative abundance of various predicted functional pathways from both the core and satellite taxa was significantly different across groups (*P <* 0.05).

**Conclusions:** Overall, we demonstrate that gut microbiota structure and predicted function differs between not only healthy controls and CP, including across CP subtypes themselves. Various taxa with important physiological roles were implicated across CP. Differences in predicted functionality of the core and satellite taxa were evident, relating to microbial processes that may have downstream consequences at the site of the GI tract. Future studies will employ larger cohorts with the incorporation of additional omics-based approaches. This will better elucidate differences in microbiota structure and functionality across CP, including relationships with pancreatic-based clinical outcomes.

## Background

Chronic pancreatitis (CP) is a multifaceted and irreversible inflammatory condition that impacts both the endocrine and exocrine functions of the pancreas (1). A common challenge in gastroenterology, its protracted course is fraught with a gamut of long-term complications (2). Diverse triggers for pancreatitis have been documented in the scientific literature, including alcohol consumption, gallstone disease, genetic mutations, metabolic disorders, iatrogenic factors, structural changes within the pancreas, infections, and environmental influences (3–6).

Small percentages of patients at risk of developing the disease will do so, alluding to other factors contributing to the development of pancreatitis and the clinical course of disease (7). Changes within the intestinal microbiota have been observed in patients with pancreatitis (8–11), however the relationship between this dysbiosis and pancreatitis is yet to be fully established at the level of causation or correlation. Furthermore, studies of gut microbiota outcomes comparing the various triggers of CP pancreatitis are scarce. In tandem with structural changes to the site of the pancreas itself, it is plausible that intestinal physiological changes associated with different CP subtypes could infer differential impacts upon intestinal microbiota composition and function. This includes the consumption of alcohol itself (12,13), alongside changes to local intestinal milieu physiology related to particular genetic mutations relevant to CP, such as within the *CFTR* gene (14). These differences are likely compounded by additional pancreatitis-related treatment interventions, such as antibiotic administration (11).

The majority of research within the CP intestinal microbiome has thus far utilised short-read amplicon sequencing across hypervariable regions of the 16S rRNA gene, due to its appeal as a cheaper high-throughput option to profile gut microbial communities (8–11,15–18). More recently, full-length 16S rRNA sequencing approaches, which offer advantageous taxonomic resolution capabilities (19), have become more economically viable for such studies as sample multiplexing capabilities continue to increase. This is particularly appealing when trying to detect potential relationships between specific bacterial species and host clinical outcomes across larger study cohorts.

In this pilot study, we employed such full-length (FL) 16S rRNA sequencing for the first time, alongside shorter amplicon (V4) sequencing, to profile microbial communities both comprehensively and with respect to current literature. Specifically, we investigated how various CP triggers, dietary (alcohol consumption) or genetic (*CFTR* mutation) in nature, might impact microbiota structure and predicted function. We further compared this data to that obtained from healthy controls for added insights. Clinical metadata was integrated to further investigate any relationships with microbiota structure and predicted function.

## Methods

### Study participants and design

CP patients from Manchester NHS Foundation Trust and healthy control volunteers were recruited between 2020 to 2022. Clinical diagnosis of CP was confirmed from hospital records and eligible participants were subsequently contacted for potential inclusion into the study. This included participants with alcohol-induced chronic pancreatitis (AIP), alongside participants genetically identified as heterozygous carriers of mutant *CFTR,* namely CFTR*-* related pancreatitis (CFRP). Healthy controls from the surrounding Manchester area were recruited. Overall, the study comprised 25 participants with CP (AIP; *n* = 11, CFRP; *n* = 14) and healthy controls (HC; *n* = 9). To account for any intra-participant temporal differences, participants were asked to donate three stool samples over a 7-day period using a home postal kit. Upon receipt, samples were stored at -80 °C at the Manchester University NHS Foundation Trust (MFT) dedicated Biobank. Basic participant clinical data was available for all groups and is summarised in Table 1. Exclusion criteria, alongside detailed clinical data for the CP groups (Table S1, is available in the Supplementary Methods and Results which are found within Additional File 1).

**Table 1.** Study group characteristics.

| Clinical variable | HC ($n = 9$) | AIP ($n = 11$) | CFRP ($n = 14$) |
| --- | --- | --- | --- |
| Age median [IQR] | 40 [26-45] | 62 [56-66] | 59 [48-66.8] |
| Body mass index median [IQR] | NC | 26.6 [22.5-30.4] | 26.8 [24.3-27.7] |
| Cholecystectomy surgery (% Yes) | 0.0 | 18.2 | 21.4 |
| Diabetes mellitus (% Yes) | 0.0 | 45.5 | 35.7 |
| Pancreatic complications (% Yes) | 0.0 | 55.5 | 14.3 |
| Pancreatic insufficiency (PI) (% Yes) | 0.0 | 81.8 | 64.3 |
| Sex (% Male) | 55.6 | 81.8 | 85.7 |
| Vitamin D ($\text{nmol}^{-1}$) median [IQR] | NC | 73.0 [43-76] | 63.8 [77-105] |

NC – not collected for respective subjects.

### PMA treatment prior to DNA extraction

1 mg propidium monoazide (PMA) (Biotium, CA, USA) was hydrated in 98 µL 20% dimethyl sulfoxide (DMSO) to give a working stock concentration of 20 mM. 300 mg of thawed stool was homogenised in 1.5 mL PBS, and then centrifuged at 3200 x g, for 5 minutes. The pellet was then resuspended in 0.5 mL PBS prior to subsequent PMA treatment. 1.25 µL of PMA (20 mM) was added to give a final concentration of 50 µM. Following the addition of PMA to samples in opaque Eppendorf tubes, PMA was mixed by vortexing for 10 seconds, followed by incubation for 15 minutes at room temperature (∼20 °C). This step was repeated before the transfer of samples to clear 1.5 mL Eppendorf tubes and placement within a LED lightbox. Treatment occurred for 15 minutes to allow PMA intercalation into DNA from compromised bacterial cells. Samples were then centrifuged at 10,000 x g for 5 minutes. The supernatant was discarded, and the cellular pellet was resuspended in 200 µL PBS. Treatment was performed to remove unviable bacteria prior to DNA extraction and subsequent sequencing, as previously described (20).

### DNA extraction process

Cellular pellets resuspended in PBS were loaded into the ZYMO Quick-DNA Fecal/Soil Microbe Miniprep Kit (Cambridge Bioscience, Cambridge, UK) as per the manufacturer’s instructions. Dual mechanical-chemical sample disruption was performed using the FastPrep-24 5G instrument (MP Biomedicals, California, USA) at a speed of 6.0 m/s for 40 seconds. Following extraction, DNA was quantity was measured using the Qubit dsDNA HS Assay Kit (high sensitivity, 0.2 to 100 ng) on the Qubit 3.0 fluorometer (Life Technologies), and purity assessed using the NanoDrop™ One Spectrophotometer (Thermo Fisher) as per the manufacturer’s protocols.

### 16S rRNA sequencing

A dual approach was undertaken to characterise the microbiota across samples. Initially, sequencing libraries targeting the V4 region of the 16S rRNA gene were prepared as per the Schloss protocol (21). Libraries were sequenced on the MiSeq using V2 2 × 250 bp chemistry. Additionally, full-length 16S rRNA gene libraries were constructed utilising the Kinnex 16S rRNA kit. These libraries were sequenced on the PacBio Revio II system. For both processes, PCR & DNA extraction negative controls were implemented, alongside the use of mock community positive controls, which included a Gut Microbiome Standard (ZYMO RESEARCH™). Sequencing was performed at the NUOmics DNA sequencing facility (Northumbria University).

### Sequence processing and bioinformatic analyses

Sequences were analysed using DADA2, as to demultiplex and remove primer sequences, validate the quality profiles of forward and reverse reads and subsequently trim, infer sequence variants, merge denoised paired-reads, remove chimeras, and finally assign taxonomy via Naive Bayesian Classifier implementation (22). Flexible assignment was implemented using a priority assignment approach: GTDB r207 (23), followed by Silva v138 (24), then lastly RefSeq + RDP (25). Remaining unidentifiable ASVs were run through BLAST (26) (https://blast.ncbi.nlm.nih.gov/Blast.cgi) and matched appropriately where possible, based on adequate query coverage. Taxa with single reads were removed and excluded from subsequent statistical analysis. Amplicon sequence variants (ASVs) from the same bacterial taxon were collapsed to form a single operational taxonomic unit (OTU) for a given taxon when applicable to the species level. The R package decontam was used to remove any potential sources of contamination across samples (27), utilising the prevalence-based contamination identification approach with a threshold classification of *P =* 0.1. To compare the predicted functional and metabolic profiles between groups, full-length 16S rRNA ASVs were analysed using PiCRUSt2 to determine Kyoto Encyclopaedia of Genes and Genomes (KEGG) orthologs (KO) and Enzyme Commission (EC) numbers (28). To provide a more comprehensive understanding of microbial functional potential, identified KOs were mapped to KEGG level 3 hierarchical pathways. Additionally, identified EC numbers were linked to predicted functions. Results from both approaches were aggregated across samples for group-wise comparisons. More detailed information on these processes is available in the Supplementary Methods and Results (Additional File 1). The dataset supporting the conclusions of this article are available within the European Nucleotide Archive repository, identifiable using accession number PRJEB78535.

### Statistical analysis

Linear regression analysis, including calculated coefficients of determination (*R*^2^), degrees of freedom (*df*), F-statistic (*F*) and significance values (*p*), were utilised for microbial partitioning into common core and rarer satellite groups, and were calculated using XLSTAT v2021.1.1 (Addinsoft, Paris, France). Fisher’s alpha index of diversity and the Bray-Curtis index of similarity were calculated using PAST v4.03 (29). Tests for significant differences in microbiota diversity were performed using Kruskal-Wallis in XLSTAT. Analysis of similarities (ANOSIM) with Bonferroni correction was used to test for significant differences in microbiota composition and functional profile and was performed in PAST. Similarity of percentages (SIMPER) analysis, to determine which constituents drove compositional differences between groups, was also performed in PAST. Ordination approaches, including stepwise selection for both canonical correspondence (CCA) and redundancy analysis (RDA), was performed in CANOCO v5 (30). Compositional data for predicted metagenome function across samples was centred log-ratio (CLR) transformed prior to heatmap generation for visualisation of sample differences across groups with ANOVA testing implemented for such comparisons. Statistical significance for all tests was deemed at the *p* ≤ 0.05 level. Subsequent figures and plots were generated using PAST (v4.03) and ggplot2 in R (v4.3.1).

## Results

Following sequence processing, FL and V4 amplicon reads of the 16S yielded similar raw counts (see Supplementary Results). Initially, we sought to partition both datasets from the whole microbiota, into the more common (≥ 75% prevalence) core taxa and the rarer, infrequent (< 75% prevalence) satellite taxa based on distribution-abundance relationships across the three study groups (Fig. 1). As such, these partitions were utilised within subsequent analyses to understand differences between microbial subcommunities and relationships with clinical outcomes across participants. Similar trends were observed across both datasets, in which the core taxa constituted 74.59% and 73.58% of the total abundance within the healthy control samples across FL and V4 datasets respectively. The satellite taxa across both AIP and CFRP were more prominent, featuring a combined relative abundance of 55.4% and 64.5% across the FL-16S, and 43.32% and 59.52% across the V4 16S rRNA datasets respectively.

**Fig. 1.**
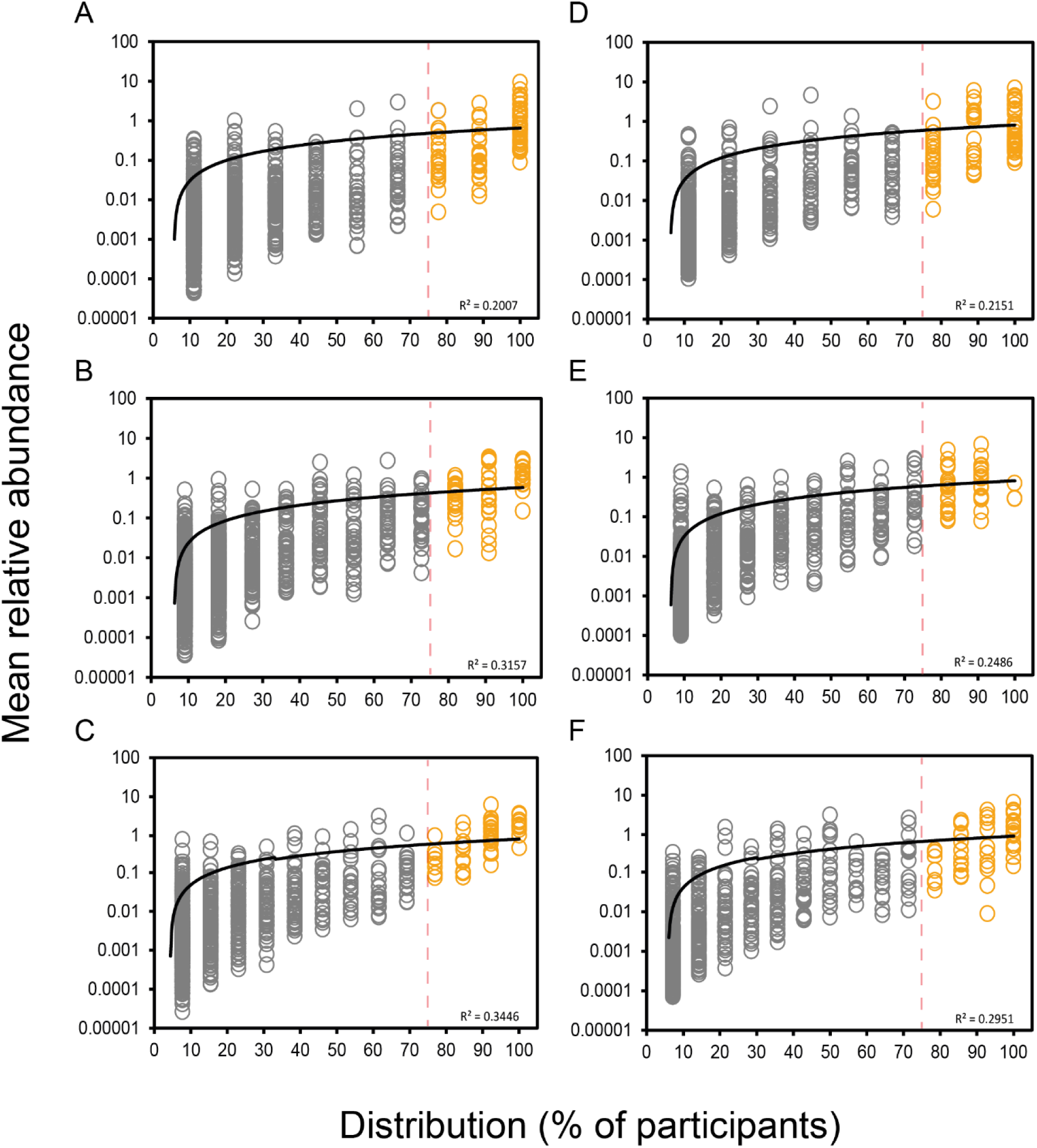
Distribution and abundance of bacterial taxa across different sample groups. **(A)** Healthy control (FL-16S). **(B)** Alcohol-induced pancreatitis (FL-16S). **(C)** Cystic fibrosis-related pancreatitis (FL-16S). **(D)** Healthy control (V4-16S). **(E)** Alcohol-induced pancreatitis (V4-16S). **(F)** Cystic fibrosis-related pancreatitis (V4-16S). Given is the percentage number of patient stool samples each bacterial taxon was observed to be distributed across, plotted against the mean percentage abundance across those samples. Core taxa are defined as those that fall within the upper quartile of distribution (orange circles), and satellite taxa (grey circles) defined as those that do not, separated by the vertical line at 75% distribution and labelled respectively. Distribution-abundance relationship regression statistics: (A) R^2^ = 0.20, F_1,1175_ = 295.1 , (B) R^2^ = 0.32, F_1,1570_ = 724.3, (C) R^2^ = 0.34, F_1,1191_ = 626.2, (D) R^2^ = 0.22, F_1,611_ = 163.30, *P <* 0.0001; (E) R^2^ = 0.25, F_1,750_ = 242.69, *P <* 0.0001; (F) R^2^ = 0.30, F_1,729_ = 301.51, *P <* 0.0001.

Comparing microbial diversity across the three study groups highlighted some significant differences (Fig 2, Table S2), which varied slightly based on the sequencing approach investigated. For the FL 16S rRNA analysis, there were no overall differences between groups in general (Fig 2A, *P =* 0.081), but pairwise differences were apparent between the AIP and CFRP groups (Fig 2A, *P =* 0.041). Core taxa diversity was more similar between groups (Fig 2A, *P =* 0.55) with no specific pairwise differences. As for the satellite taxa, again no significant overall differences between the three groups emerged (Fig 2A, *P =* 0.080). There were, however, similar differences between the AIP and CFRP groups as demonstrated prior in the whole microbiota analysis (Fig 2A, *P =* 0.041).

**Fig. 2.**
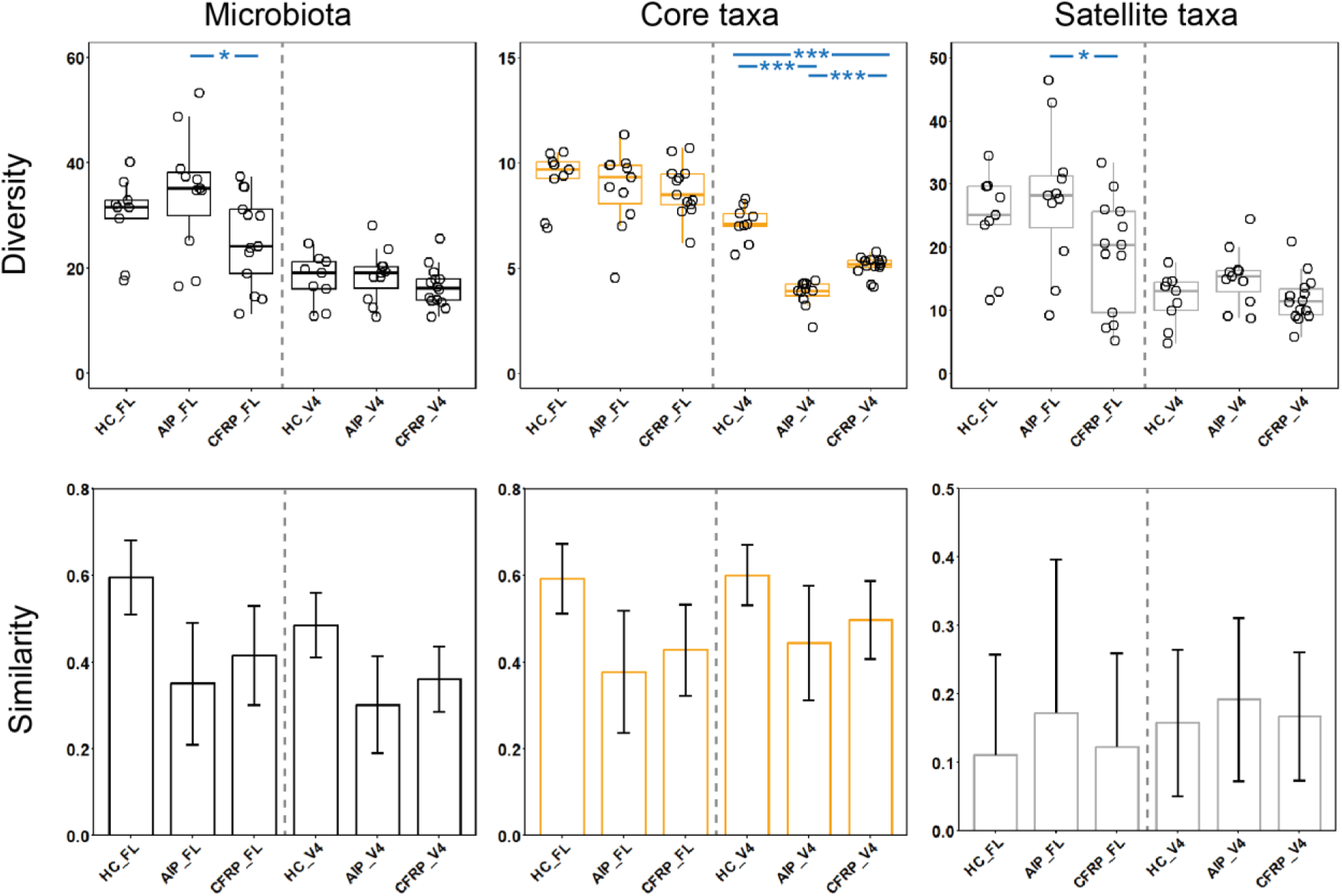
Microbiome diversity and similarity compared across healthy controls, alcohol-induced pancreatitis, and cystic fibrosis-related pancreatitis sample groups. Shown is the data for both shorter amplicon (V4) and full-length 16S rRNA sequencing data. Whole microbiota (black plots) and partitioned data into core (orange plots) and satellite taxa (grey plots) are given. **(A)** Differences in Fisher’s alpha index of diversity across groups. Black circles indicate individual patient data. Error bars represent samples that fall within the range, i.e. 1.5 times the inter-quartile range (IQR). Labelled p values describe the overall Kruskal-Wallis test values. Asterisks between groups denote a significant difference in diversity following Mann-Whitney U testing (***; p ≤ 0.0001, **; p ≤ 0.001, *; p ≤ 0.05). Summary statistics are provided in Table S2. **(B)** Microbiome variation measured within and between sampling groups, utilising the Bray-Curtis index of similarity. Error bars represent standard deviation of the mean.

For V4 16S rRNA analysis, whole microbiota diversity was similar across the three study groups (Fig. 2A, *P =* 0.32). However, when comparing microbiota partitions, differences were apparent. In particular, the core taxa diversity was significantly different across the three groups (Fig 2A, *P <* 0.0001). This included significant differences between HC and both CP groups (*P <* 0.0001) but also between both CP groups themselves (*P <* 0.0001). Satellite taxa diversity was slightly different between groups (*P =* 0.095), primarily driven by the AIP group, for which diversity was larger than both the healthy control (*P =* 0.080) and CFRP (*P =* 0.058) groups.

Despite the differences across diversity, both sequencing approaches yielded similar compositional profiles across the various microbiota partitions. In terms of whole microbiota composition, the healthy control group had the highest intra-group similarity (Fig. 2B), irrespective of the sequencing approach analysed, and clustered more tightly than the CP groups. Whilst this trend was observed across the core taxa also, the satellite taxa similarity was highly variable across all study groups (Fig. 2B). When comparing between-group microbiota composition, differences were evident across all microbiota partitions across the FL 16S rRNA analyses (Table S3). The AIP microbiota composition was consistently different to healthy controls, including at the whole microbiota (*P =* 0.048), core taxa (*P =* 0.014), and satellite taxa (*P <* 0.0001) levels. As for the CFPR group, differences compared to AIP group were evident across both the core (*P =* 0.018) and satellite (*P =* 0.0018) taxa. Significant differences between healthy controls and CFRP were only apparent across the satellite taxa (*P <* 0.0001). Across the V4 16S rRNA analysis (Table S3), differences were more modest at the whole microbiota level. However, across both the core and satellite taxa, the healthy control group were significantly different to the AIP (*P <* 0.0001) and CFRP (*P <* 0.001 core, *P <* 0.0001 satellite) groups, respectively. Interestingly, these differences also extended to between both AIP and CFRP groups across these partitions (both *P <* 0.0001).

We next sought to investigate which taxa drove the differences observed across groups, by implementation of SIMPER analysis. Across the FL 16S data (Table S4), common trends were observed between comparison of both CP groups with the healthy control group. Across the five taxa responsible for driving most dissimilarity between all groups (Fig. 3), a reduction in the mean relative abundance of *Bifidobacterium adolescentis*, *Ruminococcus bromii* and *Faecalibacterium prausnitizii* across the CP groups was a common feature that drove such dissimilarity. Other taxa that typically increased across both CP groups relative to healthy controls, included members of the genus *Streptococcus.* When directly comparing both CP groups to one another, similar taxa were implicated alongside a reduction in other taxa, including *Agathobacter rectalis* and *Akkermansia muciniphila* in the CFRP group (Table S4). When utilising the V4 16S rRNA data for the SIMPER analysis (Table S5), similar taxa were again implicated across the differences observed amongst groups, including changes to *Bifidobacterium adolescentis*, *Ruminococcus bromii*, and *Faecalibacterium prausnitizii*. The disparity amongst Proteobacterial relative abundance, namely *Escherichia*, was increased across the V4 16S rRNA analyses.

**Fig. 3.**
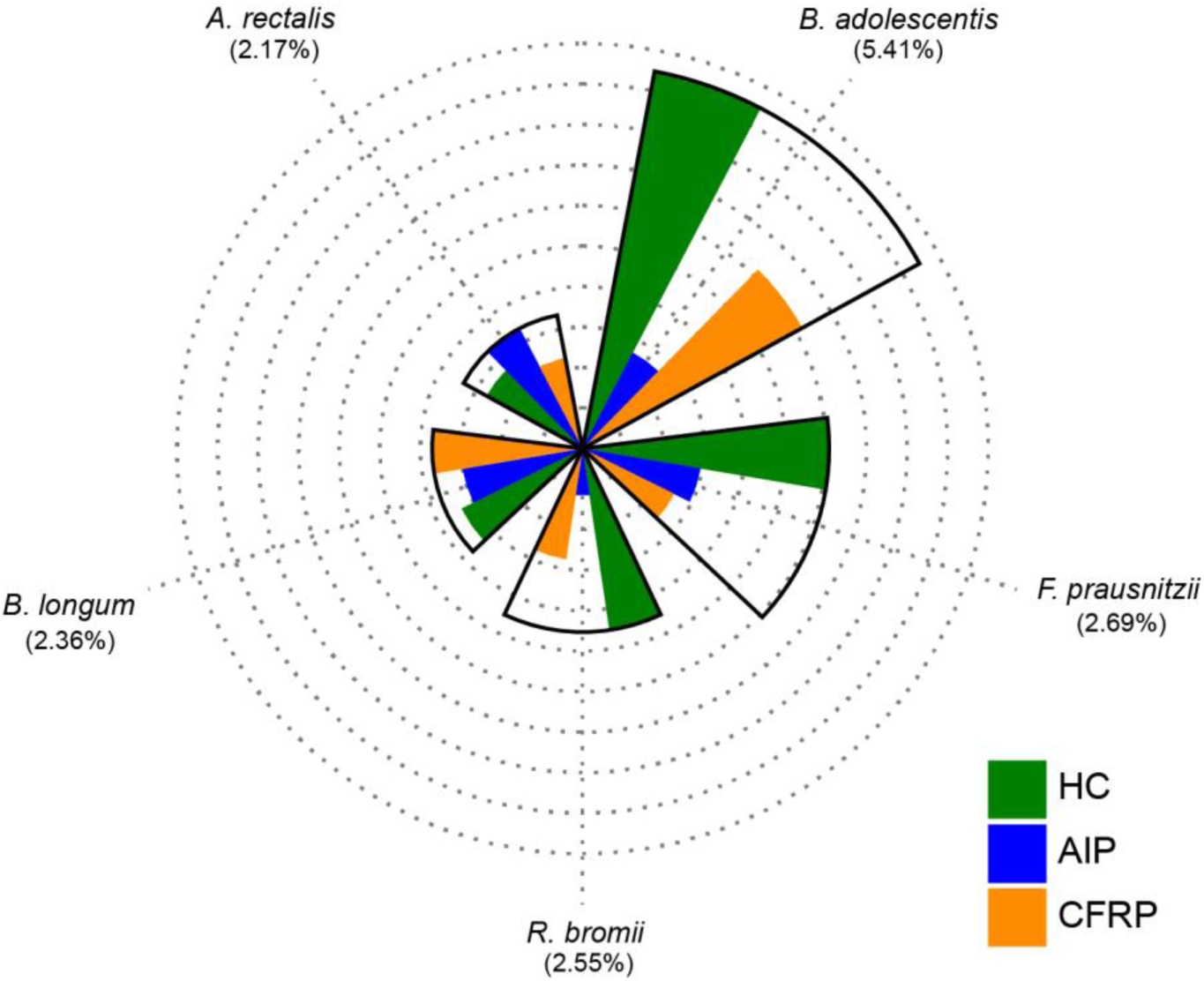
Radial bar plot depicting mean relative abundance across taxa highlighted as top drivers of dissimilarity across groups from SIMPER analysis. Each concentric ring represents an increment of 1% mean relative abundance, with the outermost ring corresponding to 10%. Average contribution to microbial dissimilarity for a given taxa across the three groups is given in parenthesis below each respective taxon. All taxa shown above were designated as core based on significant distribution-abundance relationships. Taxa are organised in clockwise fashion based on mean relative abundance across healthy controls. Detailed SIMPER results can be found in Tables S4-S5.

In addition to investigating microbiota structural differences across the study groups, we also performed analysis with PiCRUSt2 to compare predicted metagenome function, by use of associated KEGG orthologs (KOs) and enzyme classifications (ECs) across samples based on their microbiota composition. Using the FL-16S data, associated KOs across samples were mapped to KEGG hierarchical pathways to determine if differences arose between the predicted metagenome function of sampling groups. Predicted functional profile similarity was indeed highest within the healthy control group, which clustered tighter than the CP groups based on the relative predicted pathway abundance (Fig. 4). There was no defined separation overall across groups based on the 95% ellipses, though some divergence was observed among various samples from both CP groups. In terms of pathway compositional differences (Table S6), the CFRP group was more similar to healthy controls as observed across microbiota structure previously, with the AIP group being generally dissimilar to both the healthy control and CFRP groups. When comparing function inferred instead from EC counts mapped to specific KEGG modules, again similar relationships across both intra- and inter-group analyses ensued (Fig. S1).

**Fig. 4.**
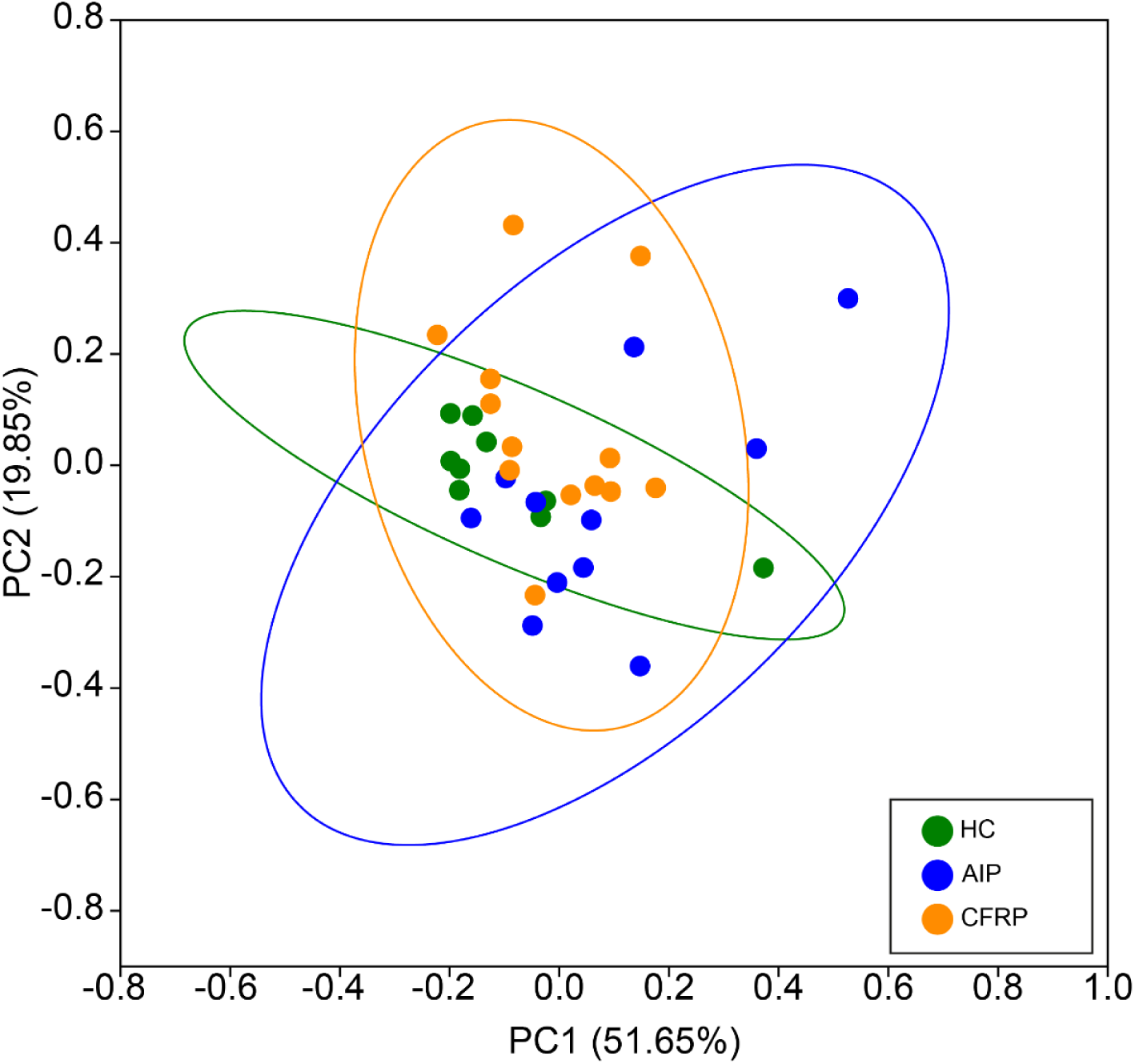
Principal Coordinates Analysis (PCoA) plot depicting compositional differences in predicted functional pathways across sample groups. Pathway abundances were derived from full-length 16S rRNA amplicon data analysed with PiCRUSt2. KEGG orthologues were aggregated prior to annotation to Level 3 hierarchical pathways. Relative abundances of pathways per sample were calculated, and Bray-Curtis dissimilarity was used as the distance metric for PCoA. ANOSIM summary statistics can be found in Table S6.

Next, participant clinical characteristics (Table 1) were incorporated into multivariate analyses to investigate potential relationships with both microbiota structure and predicted function. Across microbiota structure, CP type was a significant explanator of both the core (pseudo-*F* = 1.7, *P <* 0.008) and satellite (pseudo-*F* = 1.3, *P <* 0.004) taxa. As for predicted function, only variation within the core taxa was significantly explained by CP type, although to a greater extent (*F* = 4.1, *P <* 0.009) (Table S7).

Given the consistent impact unto both core taxa composition and predicted functional variation elicited by the type of CP present, we next compared how the most abundant predicted functions may have differed across the study groups (Fig. 5). Functions relating to key bacterial metabolic pathways were highly similar across all sampling groups. Most of the predicted functions across the core taxa were highly conserved across all groups, however there were some exceptions (Fig. 5A, Table S8). Amino sugar and nucleotide sugar metabolism was significantly different across the sampling groups (*P =* 0.008), primarily driven by differences in the AIP group compared to the healthy controls (*P =* 0.036) and CFRP groups (*P =* 0.073) respectively. As for the satellite taxa, many of these predicted associated functional pathways were highly conserved with the core taxa (Fig. 5B). There were, however, observed differences across the groups for multiple pathways (Table S8). This also included amino sugar and nucleotide sugar metabolism (*P =* 0.035) alongside ABC transporters (*P =* 0.009), quorum sensing (*P =* 0.046), starch and sucrose metabolism (*P =* 0.01), and glycolysis/gluconeogenesis-related functions (*P =* 0.025). Given that most fundamental bacterial pathways were shared across both the core and satellite taxa, we also decided to investigate how predicted pathways exclusive to the satellite taxa, which were of increased relative abundance across CP participants, may have differed between all three groups (Fig. S2, Table S8). Of the predicted level three hierarchical pathways observed, many were ubiquitously present across all study groups and mapped to the site of the GI tract. In particular, this included biosynthesis of various secondary metabolites, polycyclic aromatic hydrocarbon degradation, non-alcoholic fatty liver disease, alcoholism, and colorectal cancer.

**Fig. 5.**
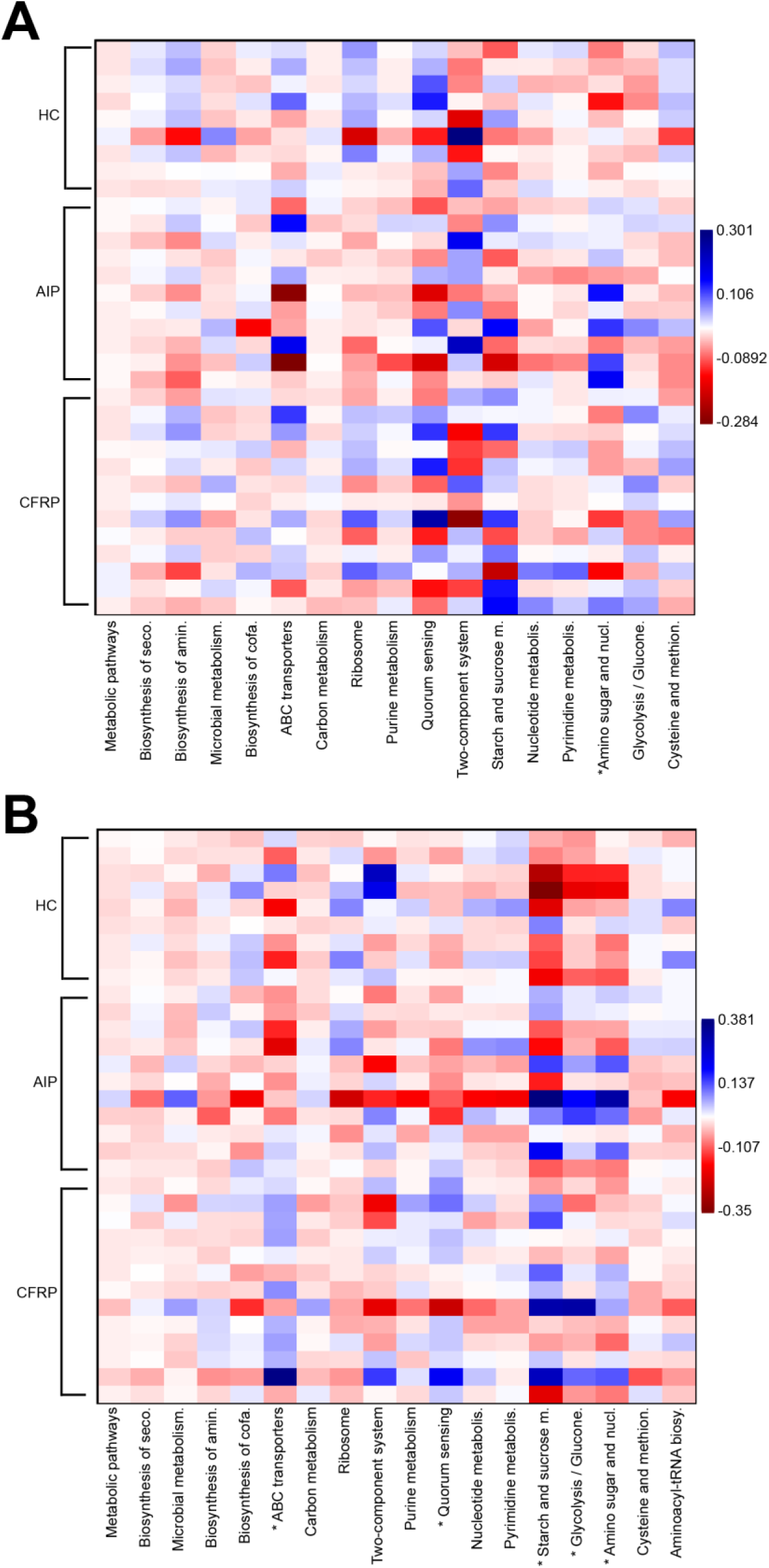
Differences in predicted metagenome function between study groups. CLR transformed compositional data was plotted to visualise differences among groups across KEGG level 3 hierarchical pathways accounting for >1% total predicted functionality. **(A)** Core taxa. **(B)** Satellite taxa. Asterisks (*) denote significant differences across group means following ANOVA testing (*P <* 0.05). Full length pathway names and associated summary statistics can be found in Table S9.

Among these pathways, only colorectal cancer was significantly different between groups (*P =* 0.047), driven by differences between the AIP and HC groups (*P =* 0.029).

## Discussion

The principal aim of this study was to investigate how intestinal microbial communities may differ among CP subtypes, including additional comparisons back to healthy controls for further interpretation. For this purpose, we piloted both full-length (FL) and shorter amplicon (V4) sequencing of the 16S rRNA gene to profile faecal microbial communities, utilising collapsed ASVs for this purpose across both approaches (31). Across similar sampling depths, as demonstrated by the raw read counts per sample post-sequencing, both approaches were comparable in partitioning total microbial communities (whole microbiota) based on significant distribution-abundance relationships across all three study groups. This is a necessary step as to isolate the common core constituents, that typically contribute a greater amount to principal functionality (in this case relating to the host), and the rarer satellite taxa which are often more opportunistic and responsive to various perturbation events across a given community (32,33). Indeed, this previously has been used across varying microbiome sites in health and disease to successfully partition taxa (34–36). Following partitioning of the microbiota into these components, it was immediately apparent that both CP groups had an expansion to the relative abundance of the satellite taxa, indicative of perturbation to the local site or changes in local physiology (37).

Irrespective of microbiota partition studied, the FL 16S rRNA data yielded larger values of diversity as compared to V4 16S approach. There were subtle similarities across both the whole microbiota and satellite taxa analyses when groups were compared, however the core taxa diversity exhibited notable differences. Whilst core taxa diversity was increased in the healthy control groups as compared to both CP groups, this was significantly pronounced in the V4 16S rRNA approach. It is plausible that enhanced species-level classification across the FL 16S rRNA data is responsible for driving such discrepancies. Various constituents of the core intestinal microbiota community, such as members of the Actinobacteria (*Bifidobacterium*) are poorly resolved solely across the V4 region (38) and may otherwise have been labelled as the same taxon. The increased diversity across FL-16S data could have also been further inflated by increased identification of single nucleotide polymorphisms across chromosomes from individual bacteria (39). This is likely to impede the satellite taxa specifically, whereby taxonomic resolution is likely hindered more than the core community, and for which clustering may not be possible at the species or genus level for a larger variety of taxa. Indeed, the disparity between diversity was largest across the satellite taxa between both sequencing approaches. Across the wider literature, there is evidence for both a reduction to gut microbiota diversity in CP (9,11,17), alongside no significant differences at all (16,18,40,41) as compared with controls. In those latter studies, CP subtypes were often grouped together for such analyses, or not explicitly depicted for sub-type comparisons. Interestingly, Zhou and colleagues showed that despite differences to controls, CP subtypes (alcohol and idiopathic) were not significantly different in terms of alpha-diversity metrics across a Chinese population (9). It remains to be seen whether partitioning of microbial communities across such participants of different geographical origin and lifestyle renders significant differences in diversity.

Despite the preliminary differences across microbial structure from the diversity observed within the current study, analyses of microbial compositions exhibited high concordance between both sequencing approaches tested, importantly validating the reliability and accuracy of such principal observations across the microbial communities. Here, the healthy control subjects had increased similarity across the whole microbiota and core taxa composition as compared to both CP groups. The increased similarity observed amongst ‘healthy’ microbial ecosystems relative to their diseased counterparts has previously been termed the ‘Anna Karenina principle’, in which dysbiotic environments can stem from a variety of different perturbation events or factors associated with a given disease (42,43). Currently, this finding is mirrored across some studies from the CP microbiota (10,17), with others indicating no increased similarity across a given group (9,11). Partitioning of microbiota into sub-communities is a pragmatic approach to decipher such relationships moving forward, given the high intragroup variance of satellite taxa composition, which seemingly drove the majority of biomass across CP participants here as observed across other dysbiotic intestinal environments (34,35,37,44). When partitioned, the composition of both core and satellite taxa across CP groups was significantly different to that of healthy controls, mirroring more general findings often observed across CP (9,11,15–18). We also demonstrated significant microbiota compositional differences between CP groups themselves across both sequencing approaches, driven by the disparity among AIP participants. Differences across disease aetiology are not established, with the current limited evidence currently available (9) conflicting across relatively low group sample sizes. From our multivariate analyses, we also did not detect any changes resultant of pancreatic complications unto the microbial communities, namely resultant of exocrine pancreatic insufficiency nor endocrine pancreatic insufficiency. Whilst both independently associate with particular microbiota structure and function (45,46), their compounding effects upon the CP microbiota are not ubiquitously reported (9–11,40).

When comparing which taxa drove the differences between sampling groups from our SIMPER analysis, various taxa from both CP groups were consistently reduced in mean relative abundance as compared to the healthy controls. This decrease was most pronounced for *Bifidobacterium adolescentis.* Members of the genus *Bifidobacterium* can exhibit anti-inflammatory effects at the site of the intestinal tract through direct modulation (47), but also through lactate production which further extends to the site of the pancreas itself (48,49). Additionally, prominent short chain fatty acid (SCFA) producers such as *Faecalibacterium prausnitzii*, alongside key intermediate fermenters, namely *Ruminococus bromii* were decreased across CP. Collectively, these findings are consistent with the wider literature, in which key members of the Firmicutes and Actinobacteria relating to maintenance of intestinal immune homeostasis are reduced in CP (9–11,15,17,18,40). It is well characterised that SCFAs, particularly butyrate, are important mediators of intestinal health relating to the modulation of local inflammatory responses and the maintenance of gut barrier integrity (50–52). On the contrary, we also observed increases in the relative abundance of *Streptococci* across CP groups. Typically designated as satellite taxa across our CP groups, *Streptococci* have previously been associated with a proinflammatory intestinal environments (53), and their relative abundance has even been suggested as a potential early biomarker for pancreatic cancers (54). These implications remain to be fully elucidated given the shorter 16S rRNA regions utilised, resulting in reduced species resolution across such studies.

An expansion of Proteobacteria across CP is well defined in the literature (8,11,15,17,55), however genera such as *Escherichia* and *Klebsiella* contributed relatively little to microbial dissimilarity between groups in our FL 16S rRNA analysis. They did, however, contribute significantly more when the V4 data was used for this purpose. This could reflect DNA library preparation biases for their detection with this primer set, or bias against them when V1-V9 (FL 16S) primers are utilised given differences observed in the GC% content across such regions (56). Other bacterial relationships were generally well maintained at the genus level between the two sequencing approaches. When comparing which taxa drove differences between the two CP groups, both sequencing approaches highlighted members of genera previously discussed, with no clear trend with respect to the mean relative abundances across these seemingly beneficial taxa. Additionally, *Akkermansia muciniphila*, which reportedly exerts beneficial protective effects upon intestinal inflammation and other gastrointestinal abnormalities (57,58), was shown to be decreased across the CFRP group as compared to the AIP participants. Decreases in *A. muciniphila* are not widely reported across human CP microbiota studies yet feature commonly within cystic fibrosis intestinal microbiota dysbiosis (59–61). Given its utilisation of mucin as a primary source of nutrients, disruption to mucosal viscosity from heterozygous carriage of mutant *CFTR* and subsequent decrease in extracellular chloride and bicarbonate secretion could in theory exacerbate any diminishing effects upon *A. muciniphila* relative abundance in CP. Murine studies have further demonstrated bidirectionality between *A. muciniphila* abundance and optimal pancreatic outcomes (62,63).

Following the high overall concordance of relationships between groups across both the FL and V4 16S rRNA compositional and SIMPER tests, we confidently opted to prioritise the FL data for subsequent analyses, including predictive metagenome functionality. Modest differences were highlighted across the study groups. The healthy control samples clustered tightly to one another, whilst the two CP groups demonstrated higher intra-sample variation. Between groups, the differences between the healthy control and AIP groups were most pronounced. For comparative purposes and subsequent analyses, we utilised annotated KEGG pathways at the level 3 hierarchy, given the increased detail of biochemical pathways, versus the more general functions and broader categories exhibited at lower levels. This was likely to ascertain any meaningful differences between groups, given that high similarity and functional redundancy is readily observed across individuals with different microbiota structures (64,65). Indeed, when comparing KEGG level 3 hierarchical pathways between groups, many key components of bacterial function and metabolism were highly conserved across not only the sampling groups, but across both partitions of the microbiota itself.

Of the most abundant pathways (%) identified from the core taxa overall, only amino and nucleotide sugar metabolism was significantly different across CP, which was increased in the AIP group specifically. Similarly increased in AIP amongst the satellite taxa, these implications remain to be further explored but may indicate adaptation of the community to utilise available substrates more effectively in the presence of environmental changes or stressors to local intestinal physiology (66). Other differences among the satellite taxa were pathways such as those related to ABC transporters (increased in CFRP). ABC transporters are a diverse class of membrane proteins, with key roles in virulence across prokaryotes, which involves the uptake of essential nutrients, alongside efflux of toxins and other substances to their local environment to enhance their survival and overall fitness (67,68). They have previously been shown as elements of faecal metagenomic signatures relating to pancreatic cancer (69). The CFRP group also demonstrated increases to pathways related to quorum sensing, which might further underpin effective expansion and virulence factor coordination across the satellite taxa in the presence of dysmotility and general bacterial overgrowth in CP (70,71). The potential importance of satellite was finally highlighted through our analysis of associated functional pathways exclusively found across this subcommunity. Of the GI-related pathways exclusively identified, an increase in significant relative enrichment of pathways relating to colorectal cancer could be seen across CP (particularly CFRP) participants. Further work understanding which elements of the microbiome are attributable to this KEGG analysis is warranted but likely represents a combination of both direct and indirect microbiota-driven actions that affect host physiology and promote inflammatory pathways.

These results of course remain to be validated, with wider implications explored using more sophisticated multi-omics approaches, as only a single study across the CP microbiome thus far has utilising metabolomics alongside metagenomics or 16S rRNA gene sequencing for example (18). Additionally, larger cohorts of participants with alcohol and genetic-related pancreatitis holds considerable appeal, given the observed differences observed within our pilot analyses. This will inherently allow for more robust investigations into relationships between specific pancreatic manifestations and the gut microbiome. Whilst we had extensive data surrounding pancreatic sub-complications across our cohort, including pancreatic ductal strictures, pseudocysts, and necrosis, our limited sample size prevented reliable analysis of such factors. Given the proposed gut-pancreas axis, it should be of keen interest to determine signatures of gut microbiota translocation, or their metabolites, to such pancreatic tissues across CP (72). Other lifestyle factors such as smoking status, were collected here but were exclusively co-correlated with the AIP group for the purpose of our multivariate approach. Additional exploration into smoking as a confounder may be warranted due to it impact upon CP and elements of the microbiota (11,73).

Aside from the limitations of sample size inherent to the nature of pilot studies themselves, which here was further impacted by the COVID-19 pandemic, we also acknowledge some other caveats of this study. Firstly, we understand the predictive nature of microbiota functionality should be considered putative given the sole use of the 16S rRNA gene to infer this function. For this reason, we only utilised full-length 16S rRNA sequencing, as to resolve taxa more confidently down to the species level and use such ASVs for subsequent bioinformatic pipelines. Nonetheless, the sole use of shorter hypervariable 16S regions have previously been used to infer microbial function across the gut microbiome in CP and pancreatic cancer (10,16–18). Whilst our inclusion criteria specifically excluded participants with recent antibiotic usage, we understand that antibiotic administration across CP patients to treat infections associated with the aforementioned sub-complications (74,75) are likely historical perturbation events within our older CP cohort. It may be of interest to retrospectively investigate antibiotic usage across participants moving forward, or rather investigate microbiome dynamics longitudinally with increased granularity, as to determine the impacts of such events upon intestinal microbial communities. Whilst our smaller sample size does limit statistical power for extensive investigations, it also highlights the principal strength of this study, which is its ability to highlight microbiota differences across CP subtypes based on both structure and predicted metagenome functions. This warrants additional investigation into more specific relationships across GI physiology and the microbiota in CP.

## Conclusions

Overall, we implemented full length 16S rRNA sequencing for the first time to demonstrate that gut microbiota structure and predicted function differs between not only healthy controls and CP, but across CP subtypes themselves. This may be in part due to inherent differences across intestinal and pancreatic physiology, alongside lifestyle factors relating to disease. Differences in predicted functionality of the core and satellite taxa are evident, relating to microbial processes that may have downstream physiological consequences at the site of the GI tract. Future studies will employ larger cohorts with the incorporation of additional omics-based approaches. This will better elucidate differences in microbiota structure and functionality across CP subtypes, including relationships with pancreatic-based clinical outcomes.

## List of Abbreviations

CP: Chronic Pancreatitis
AIP: Alcohol-induced CP
CFRP: Cystic fibrosis-related CP
HC: Healthy controls
GI: Gastrointestinal
PMA: Propidium monoazide
DMSO: Dimethyl sulfoxide
FL: Full-length (16S)
ASV: Amplicon Sequence Variant
OTU: Operational Taxonomic Unit
KEGG: Kyoto Encyclopedia of Genes and Genomes
KO: KEGG orthologue
EC: Enzyme commission
CCA: Canonical correlation analysis
RDA: Redundancy analysis
CLR: Centred log-ratio transformation
SIMPER: Similarity of percentages ANOSIM Analysis of similarity
PCoA: Principal co-ordinate analysis

## Supplementary Information

Supplementary Methods and Results are available in Additional file 1.

## Acknowledgements

The authors would like to thank all participants for their involvement in the study. We also thank technical staff at the Manchester University NHS Foundation Trust biobank, for their help with sample collection and storage.

## Authors’ contributions

AYA, DWR, JM, CvdG, RM contributed to the study conception and design, patient recruitment, sample collection, data collection and interpretation. AYA, AR, CF, DWR, CvdG, RM contributed to the bioinformatics, statistical analysis, and interpretation of results. AYA, CvdG, RM contributed to the original draft of the manuscript. All authors contributed to the final draft of the manuscript.

## Data availability

The sequencing datasets generated for the current study are available in the European Nucleotide Archive (ENA) repository (https://www.ebi.ac.uk/ena/browser/view/PRJEB78535), under the title of “A cross-sectional pilot study investigating the microbiota across participants with varying types of chronic pancreatitis (CP) & healthy controls” with accession number PRJEB78535.

## Declarations

## Ethical approval and consent to participate

The current study was conducted in accordance with the principles of the Declaration of Helsinki. Ethical approval was obtained from NHS East Midlands-Nottingham Research Ethics Committee under REC reference 20/EM/0016. All participants provided signed written informed consent.

## Consent for publication

Not applicable.

## Additional File 1 – Supplementary Methods & Results

### Supplementary Methods

#### Study participants and design

Exclusion criteria for all participants included abstinence from prior antibiotic use within the previous 3 months, no diagnosis of alternate gastrointestinal conditions, abstinence from illicit drug use, and finally having no prior experience of diarrhoea lasting in excess of 48 hours within the last 6 months. All participants were asked to donate three stool samples using a home postal kit over the course of a 7-day period, which was met with relatively high adherence of 2.77 ± 0.55 (mean ± SD), to account for any intra-individual temporal variance. Upon receipt, samples were immediately stored at -80 °C at the Manchester Foundation Trust dedicated Biobank. Samples were individually processed and sequenced across both platforms. Additionally, basic participant clinical data was available for all groups, and is summarised in Table 1. Additional metadata, including specific pancreatic complications, surgical and endoscopic interventions, were reported by the consultant gastroenterologists following routine clinical practices.

#### Predictive functional annotation using PICRUSt2

To provide a more comprehensive understanding of microbial functional potential, identified KEGG orthologues (KO) were mapped to level 3 identified pathway. Additionally, identified enzyme commission (EC) numbers were linked to predicted functions. Results from both approaches were aggregated across samples for group-wise comparisons. This mapping was based on the KEGG pathway maps br08901 of BRITE Functional Hierarchies in the KEGG database (http://www.genome.jp/kegg-bin/get_htext?br08901.keg). The classification of enzyme functions was based on KEGG pathway map01100.keg from the BRITE Functional Hierarchies.

### Supplementary Results

Following sequencing processing of the 16S rRNA gene data obtained from the Illumina MiSeq, a total of 4,704,612 reads were obtained, averaging (± SD) 49,522 ± 7,964 reads per sample. Full-length 16S rRNA sequencing on the PacBio Revio II system yielded similar values, whereby a total of 3,123,278 reads were obtained, averaging (± SD) 36,744 ± 26,768 reads per sample. Samples with excessively low reads (<1000, n = 1) were removed from subsequent analyses due to unsuitability for subsequent similarity and diversity analyses.

**Table S1.** Individual metadata across the CP cohort.

| ID | Group | Endocrine<br>insufficiency | Exocrine<br>insufficiency | PERT | Endoscopic<br>intervention | Cholecystectomy | Pancreatic<br>complications | Smoker | Vitamin D<br>(nmol) |
| --- | --- | --- | --- | --- | --- | --- | --- | --- | --- |
| 016 | AIP | N | Y | Y | AD, ERCP (fistula closure) | N | PN | Y | 9.3 |
| 015 | AIP | Y | Y | Y | N | N | N | N | - |
| 017 | AIP | Y | Y | Y | ERCP (CBD stricture) | N | N | Y | 76.0 |
| 019 | AIP | Y | Y | Y | AD | N | PC | Y | - |
| 023 | AIP | N | Y | Y | AD | N | PC | Y | 73.0 |
| 018 | AIP | N | Y | Y | AD | N | PN | X | - |
| 020 | AIP | Y | Y | Y | None | N | N | N | - |
| 033 | AIP | Y | Y | Y | CB | N | N | Y | - |
| 031 | AIP | N | N | N | N | Y | PC | X | - |
| 028 | AIP | N | Y | Y | AD | Y | PN | Y | 117.0 |
| 030 | AIP | N | N | N | N | N | N | Y | 43.0 |
| 012 | CFRP | Y | Y | Y | ERCP | Y | CBD stones | N | - |
| 006 | CFRP | N | N | Y | N | N | N | N | 146.0 |
| 026 | CFRP | N | Y | Y | N | N | N | N | 113.0 |
| 034 | CFRP | N | Y | Y | N | N | N | N | 18.9 |
| 014 | CFRP | N | Y | Y | N | N | PN, PA | N | 67.8 |
| 008 | CFRP | N | N | N | N | N | N | N | - |
| 001 | CFRP | Y | Y | Y | N | N | N | X | 106.0 |
| 003 | CFRP | N | Y | Y | N | N | N | N | - |
| 007 | CFRP | N | N | N | N | Y | N | N | 72.0 |
| 010 | CFRP | Y | Y | Y | ERCP, DS | N | PDS | X | 62.5 |
| 011 | CFRP | N | N | N | N | N | N | N | - |
| 013 | CFRP | Y | Y | Y | N | N | N | N | 82.0 |
| 004 | CFRP | N | N | N | N | N | N | X | 16.0 |
| 009 | CFRP | Y | Y | Y | ERCP | Y | CBD stones | N | 102.0 |

AIP; Alcohol-induced pancreatitis, AD; Axios drainage, CB; Coelic block, CBD; Common bile duct, CFRP; CFTR-related pancreatitis, DS; Pancreatic duct stent, ECRP; Endoscopic retrograde cholangiopancreatography, PA; Pseudoaneurysm, PC; Pseudocyst, PDS; Pancreatic distal stricture, PN; Pancreas necrosis, X; Ex-smoker. Y; Yes, N; No. Absent data is denoted with '-'.

**Table S2.**
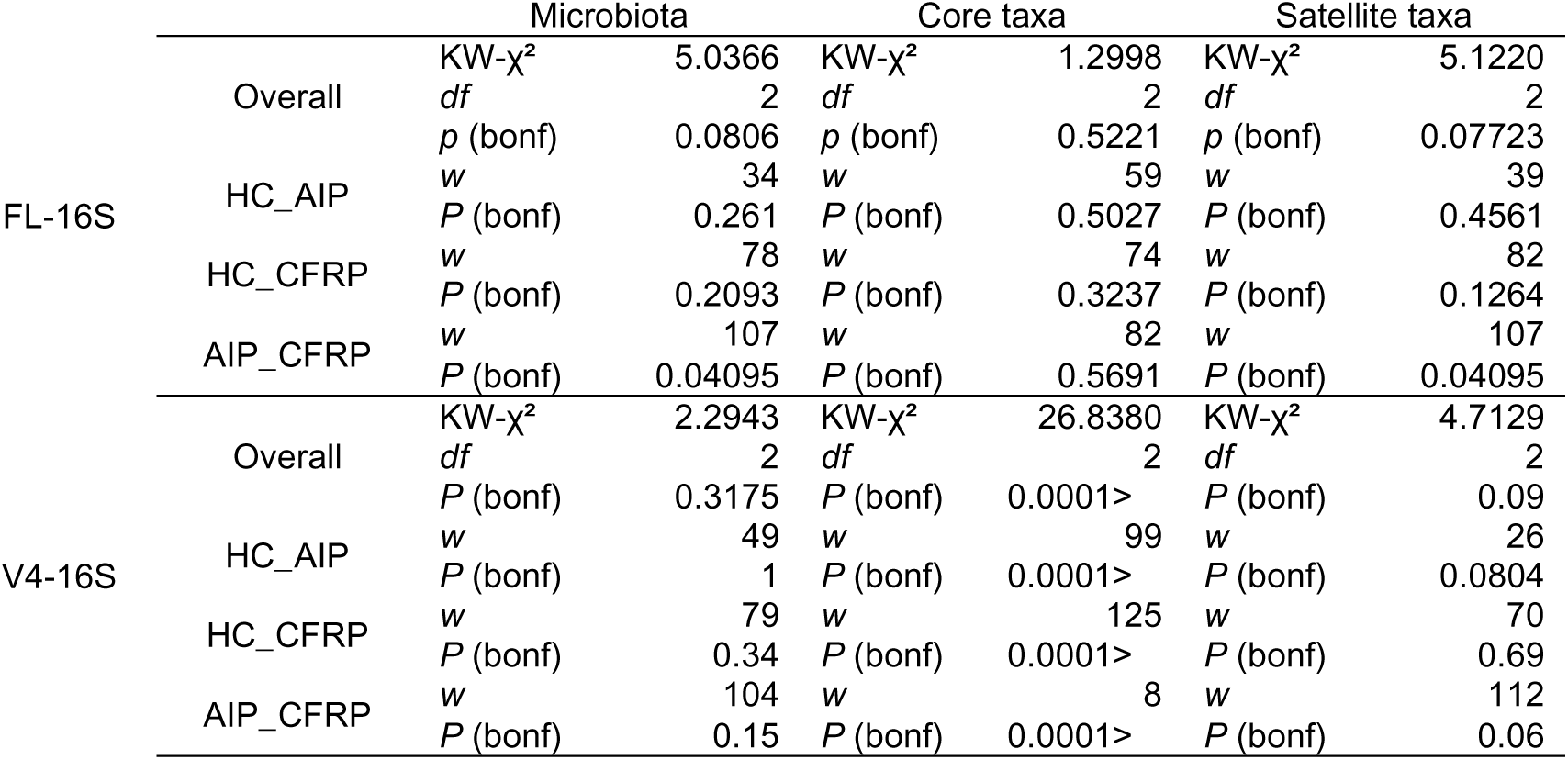
Microbiota diversity summary statistics between groups across various microbiota partitions.

**Table S3.** ANOSIM summary statistics across both FL and V4 16S rRNA analyses of microbiota composition.

|  |  | Microbiota |  | Core taxa |  | Satellite taxa |  |
| --- | --- | --- | --- | --- | --- | --- | --- |
| FL-16S | HC_AIP | *R* | 0.1011 | *R* | 0.1628 | *R* | 0.4332 |
|  |  | *P (same)* | 0.0483 | *P (same)* | 0.0143 | *P (same)* | 0.0001 |
|  |  | *P (bonf)* | 0.0489 | *P (bonf)* | 0.0151 | *P (bonf)* | 0.0002 |
|  |  | Perm N | 9999 | Perm N | 9999 | Perm N | 9999 |
|  | HC_CFRP | *R* | -0.01335 | *R* | 0.05083 | *R* | 0.3321 |
|  |  | *P (same)* | 0.523 | *P (same)* | 0.207 | *P (same)* | 0.0001 |
|  |  | *P (bonf)* | 0.5272 | *P (bonf)* | 0.2084 | *P (bonf)* | 0.0002 |
|  |  | Perm N | 9999 | Perm N | 9999 | Perm N | 9999 |
|  | AIP_CFRP | *R* | 0.0499 | *R* | 0.1262 | *R* | 0.1984 |
|  |  | *P (same)* | 0.1343 | *P (same)* | 0.0181 | *P (same)* | 0.0018 |
|  |  | *P (bonf)* | 0.1369 | *P (bonf)* | 0.0187 | *P (bonf)* | 0.0028 |
|  |  | Perm N | 9999 | Perm N | 9999 | Perm N | 9999 |
| V4-16S | HC_AIP | *R* | 0.04762 | *R* | 0.5376 | *R* | 0.5553 |
|  |  | *P (same)* | 0.1628 | *P (same)* | 0.0001 | *P (same)* | 0.0001 |
|  |  | *P (bonf)* | 0.1666 | *P (bonf)* | 0.0001 | *P (bonf)* | 0.0001 |
|  |  | Perm N | 9999 | Perm N | 9999 | Perm N | 9999 |
|  | HC_CFRP | *R* | -0.06149 | *R* | 0.3441 | *R* | 0.5572 |
|  |  | *P (same)* | 0.782 | *P (same)* | 0.0002 | *P (same)* | 0.0001 |
|  |  | *P (bonf)* | 0.791 | *P (bonf)* | 0.0002 | *P (bonf)* | 0.0001 |
|  |  | Perm N | 9999 | Perm N | 9999 | Perm N | 9999 |
|  | AIP_CFRP | *R* | 0.01308 | *R* | 0.4217 | *R* | 0.2988 |
|  |  | *P (same)* | 0.3506 | *P (same)* | 0.0001 | *P (same)* | 0.0001 |
|  |  | *P (bonf)* | 0.3534 | *P (bonf)* | 0.0001 | *P (bonf)* | 0.0001 |
|  |  | Perm N | 9999 | Perm N | 9999 | Perm N | 9999 |

**Table S4.**
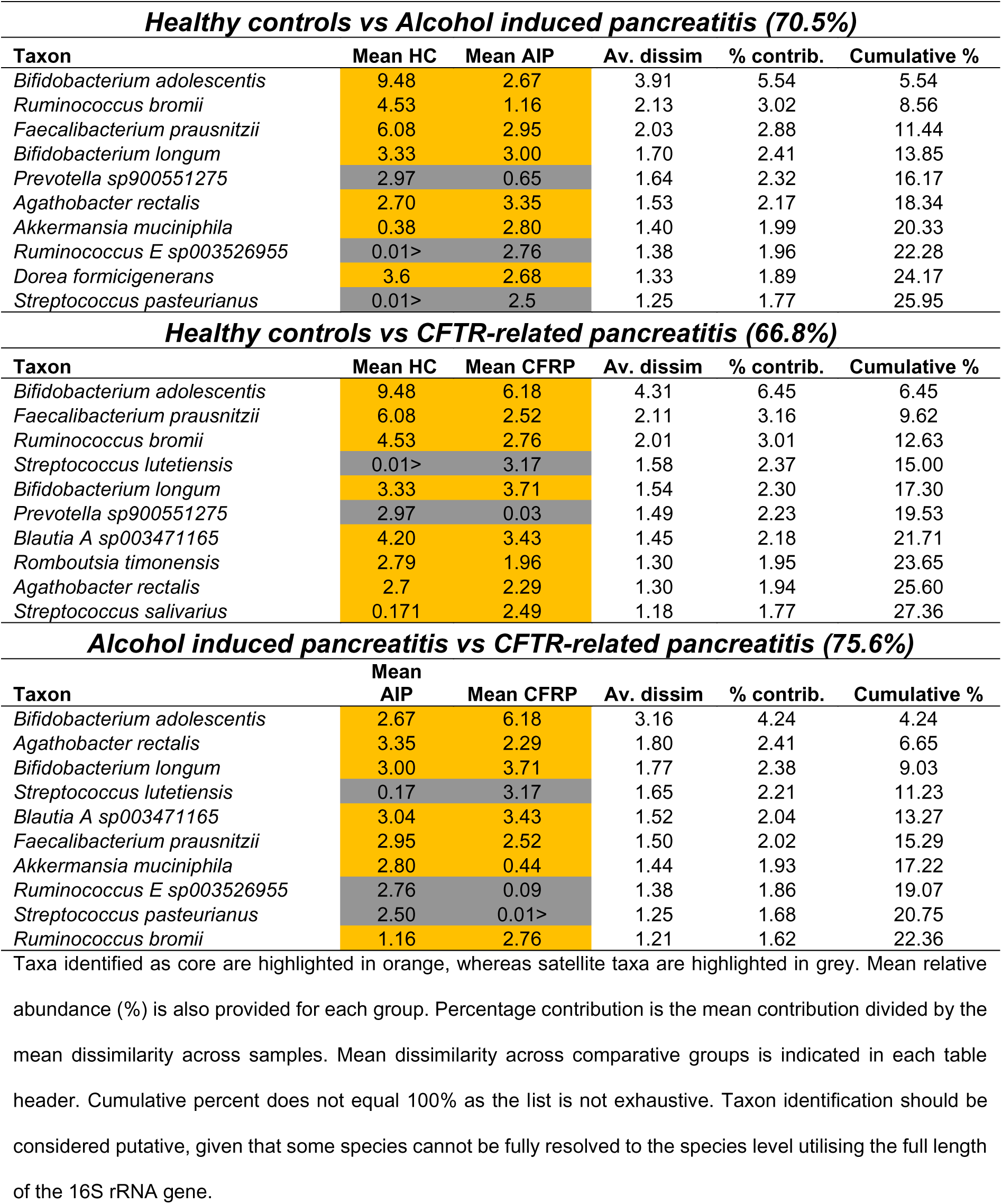
Similarity of percentage (SIMPER) analysis of microbiota dissimilarity (Bray-Curtis) between study groups, utilsing the full-length 16S rRNA sequencing data.

| **Healthy controls vs Alcohol induced pancreatitis (70.5%)** |  |  |  |  |  |
| --- | --- | --- | --- | --- | --- |
| **Taxon** | **Mean HC** | **Mean AIP** | **Av. dissim** | **% contrib.** | **Cumulative %** |
| *Bifidobacterium adolescentis* | 9.48 | 2.67 | 3.91 | 5.54 | 5.54 |
| *Ruminococcus bromii* | 4.53 | 1.16 | 2.13 | 3.02 | 8.56 |
| *Faecalibacterium prausnitzii* | 6.08 | 2.95 | 2.03 | 2.88 | 11.44 |
| *Bifidobacterium longum* | 3.33 | 3.00 | 1.70 | 2.41 | 13.85 |
| *Prevotella sp900551275* | 2.97 | 0.65 | 1.64 | 2.32 | 16.17 |
| *Agathobacter rectalis* | 2.70 | 3.35 | 1.53 | 2.17 | 18.34 |
| *Akkermansia muciniphila* | 0.38 | 2.80 | 1.40 | 1.99 | 20.33 |
| *Ruminococcus E sp003526955* | 0.01> | 2.76 | 1.38 | 1.96 | 22.28 |
| *Dorea formicigenerans* | 3.6 | 2.68 | 1.33 | 1.89 | 24.17 |
| *Streptococcus pasteurianus* | 0.01> | 2.5 | 1.25 | 1.77 | 25.95 |
| **Healthy controls vs CFTR-related pancreatitis (66.8%)** |  |  |  |  |  |
| **Taxon** | **Mean HC** | **Mean CFRP** | **Av. dissim** | **% contrib.** | **Cumulative %** |
| *Bifidobacterium adolescentis* | 9.48 | 6.18 | 4.31 | 6.45 | 6.45 |
| *Faecalibacterium prausnitzii* | 6.08 | 2.52 | 2.11 | 3.16 | 9.62 |
| *Ruminococcus bromii* | 4.53 | 2.76 | 2.01 | 3.01 | 12.63 |
| *Streptococcus lutetiensis* | 0.01> | 3.17 | 1.58 | 2.37 | 15.00 |
| *Bifidobacterium longum* | 3.33 | 3.71 | 1.54 | 2.30 | 17.30 |
| *Prevotella sp900551275* | 2.97 | 0.03 | 1.49 | 2.23 | 19.53 |
| *Blautia A sp003471165* | 4.20 | 3.43 | 1.45 | 2.18 | 21.71 |
| *Romboutsia timonensis* | 2.79 | 1.96 | 1.30 | 1.95 | 23.65 |
| *Agathobacter rectalis* | 2.7 | 2.29 | 1.30 | 1.94 | 25.60 |
| *Streptococcus salivarius* | 0.171 | 2.49 | 1.18 | 1.77 | 27.36 |
| **Alcohol induced pancreatitis vs CFTR-related pancreatitis (75.6%)** |  |  |  |  |  |
| **Taxon** | **Mean AIP** | **Mean CFRP** | **Av. dissim** | **% contrib.** | **Cumulative %** |
| *Bifidobacterium adolescentis* | 2.67 | 6.18 | 3.16 | 4.24 | 4.24 |
| *Agathobacter rectalis* | 3.35 | 2.29 | 1.80 | 2.41 | 6.65 |
| *Bifidobacterium longum* | 3.00 | 3.71 | 1.77 | 2.38 | 9.03 |
| *Streptococcus lutetiensis* | 0.17 | 3.17 | 1.65 | 2.21 | 11.23 |
| *Blautia A sp003471165* | 3.04 | 3.43 | 1.52 | 2.04 | 13.27 |
| *Faecalibacterium prausnitzii* | 2.95 | 2.52 | 1.50 | 2.02 | 15.29 |
| *Akkermansia muciniphila* | 2.80 | 0.44 | 1.44 | 1.93 | 17.22 |
| *Ruminococcus E sp003526955* | 2.76 | 0.09 | 1.38 | 1.86 | 19.07 |
| *Streptococcus pasteurianus* | 2.50 | 0.01> | 1.25 | 1.68 | 20.75 |
| *Ruminococcus bromii* | 1.16 | 2.76 | 1.21 | 1.62 | 22.36 |

Taxa identified as core are highlighted in orange, whereas satellite taxa are highlighted in grey. Mean relative abundance (%) is also provided for each group. Percentage contribution is the mean contribution divided by the mean dissimilarity across samples. Mean dissimilarity across comparative groups is indicated in each table header. Cumulative percent does not equal 100% as the list is not exhaustive. Taxon identification should be considered putative, given that some species cannot be fully resolved to the species level utilising the full length of the 16S rRNA gene.

**Table S5.** Similarity of percentage (SIMPER) analysis of microbiota dissimilarity (Bray-Curtis) between study groups, utilsing the V4 16S rRNA sequencing data.

| **Healthy controls vs Alcohol induced pancreatitis (63.5%)** |  |  |  |  |  |
| --- | --- | --- | --- | --- | --- |
| **Taxon** | **Mean HC** | **Mean AIP** | **Av. dissim** | **% Contrib.** | **Cumulative %** |
| *Escherichia coli* | 3.22 | 6.77 | 3.62 | 5.71 | 5.71 |
| *Bifidobacterium adolescentis* | 6.07 | 1.72 | 2.62 | 4.13 | 9.84 |
| *Prevotella_9 copri* | 4.60 | 1.02 | 2.51 | 3.96 | 13.79 |
| *Faecalibacterium prausnitzii* | 7.05 | 4.83 | 2.05 | 3.23 | 17.03 |
| *Ruminococcus bromii* | 4.02 | 1.58 | 1.85 | 2.91 | 19.94 |
| *Subdoligranulum variabile* | 3.18 | 2.94 | 1.52 | 2.39 | 22.33 |
| *Blautia massiliensis* | 4.48 | 2.08 | 1.42 | 2.24 | 24.56 |
| *Akkermansia muciniphila* | 0.52 | 2.81 | 1.41 | 2.23 | 26.79 |
| *Agathobacter rectalis* | 3.33 | 2.96 | 1.31 | 2.06 | 28.85 |
| *Ruminococcoides bili* | 0.00 | 2.54 | 1.27 | 2.00 | 30.85 |
| **Healthy controls vs CFTR-related pancreatitis (59.5%)** |  |  |  |  |  |
| **Taxon** | **Mean HC** | **Mean CFRP** | **Av. dissim** | **% Contrib.** | **Cumulative %** |
| *Escherichia coli* | 3.22 | 4.19 | 2.77 | 4.66 | 4.66 |
| *Bifidobacterium adolescentis* | 6.07 | 3.19 | 2.71 | 4.56 | 9.22 |
| *Prevotella_9 copri* | 4.60 | 0.16 | 2.33 | 3.91 | 13.12 |
| *Agathobacter rectalis* | 3.33 | 4.18 | 2.00 | 3.37 | 16.49 |
| *Ruminococcus bromii* | 4.02 | 2.65 | 1.82 | 3.05 | 19.54 |
| *Faecalibacterium prausnitzii* | 7.05 | 6.57 | 1.53 | 2.57 | 22.11 |
| *Subdoligranulum variabile* | 3.18 | 3.15 | 1.41 | 2.37 | 24.47 |
| *Blautia OTU 2* | 4.14 | 3.52 | 1.33 | 2.23 | 26.70 |
| *Faecalibacterium duncaniae* | 3.66 | 1.93 | 1.28 | 2.15 | 28.85 |
| *Catenibacterium mitsuokai* | 2.43 | 0.00 | 1.22 | 2.04 | 30.89 |
| **Alcohol induced pancreatitis vs CFTR-related pancreatitis (66.4%)** |  |  |  |  |  |
| **Taxon** | **Mean AIP** | **Mean CFRP** | **Av. dissim** | **% Contrib.** | **Cumulative %** |
| *Escherichia coli* | 6.77 | 4.19 | 3.79 | 5.71 | 5.71 |
| *Faecalibacterium prausnitzii* | 4.83 | 6.57 | 2.30 | 3.47 | 9.17 |
| *Agathobacter rectalis* | 2.96 | 4.18 | 2.25 | 3.39 | 12.56 |
| *Bifidobacterium adolescentis* | 1.72 | 3.19 | 1.79 | 2.69 | 15.25 |
| *Akkermansia muciniphila* | 2.81 | 1.18 | 1.65 | 2.49 | 17.73 |
| *Subdoligranulum variabile* | 2.94 | 3.15 | 1.64 | 2.47 | 20.21 |
| *Blautia OTU 2* | 3.45 | 3.52 | 1.49 | 2.24 | 22.44 |
| *Blautia massiliensis* | 2.08 | 3.97 | 1.36 | 2.04 | 24.48 |
| *Ruminococcus bromii* | 1.58 | 2.65 | 1.27 | 1.92 | 26.40 |
| *Ruminococcoides bili* | 2.54 | 0.09 | 1.27 | 1.92 | 28.31 |

Taxa identified as core are highlighted in orange, whereas satellite taxa are highlighted in grey. Mean relative abundance (%) is also provided for each group. Percentage contribution is the mean contribution divided by the mean dissimilarity across samples. Mean dissimilarity across comparative groups is indicated in each table header. Cumulative percent does not equal 100% as the list is not exhaustive. Taxon identification should be considered putative, given that some species cannot be fully resolved to the species level utilising the V4 region of the 16S rRNA gene.

**Table S6.** ANOSIM summary statistics across relative pathway abundance for annotated KEGG level 3 heirarchical pathways across all three study groups, utilising whole microbiota composition.

| Microbiota |  |  |
| --- | --- | --- |
| **HC_AIP** | *R* | 0.1116 |
|  | *P (same)* | 0.0587 |
|  | *P (bonf)* | 0.0597 |
|  | Perm N | 9999 |
| **HC_CFRP** | *R* | -0.01874 |
|  | *P (same)* | 0.5339 |
|  | *P (bonf)* | 0.5374 |
|  | Perm N | 9999 |
| **AIP_CFRP** | *R* | 0.03528 |
|  | *P (same)* | 0.2001 |
|  | *P (bonf)* | 0.2048 |
|  | Perm N | 9999 |

**Fig. S1.**
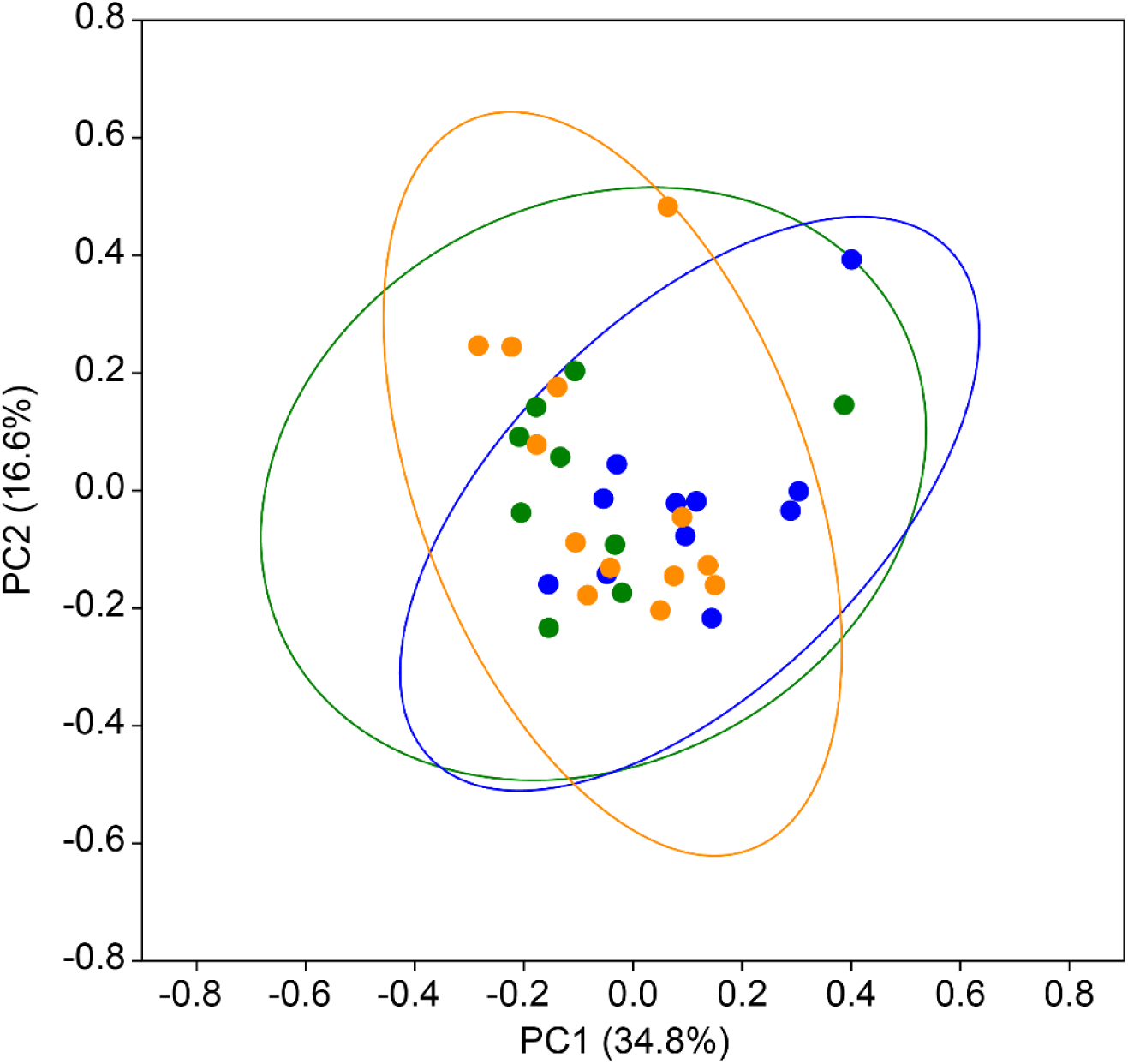
Principal Coordinates Analysis (PCoA) plot depicting compositional differences in predicted functional pathways across sample groups. Pathway abundances were derived from full-length 16S rRNA amplicon data analysed with PiCRUSt2. EC modules were aggregated prior to annotation to Level 3 hierarchical pathways. Relative abundances of pathways per sample were calculated, and Bray-Curtis dissimilarity was used as the distance metric for PCoA.

**Table S7.**
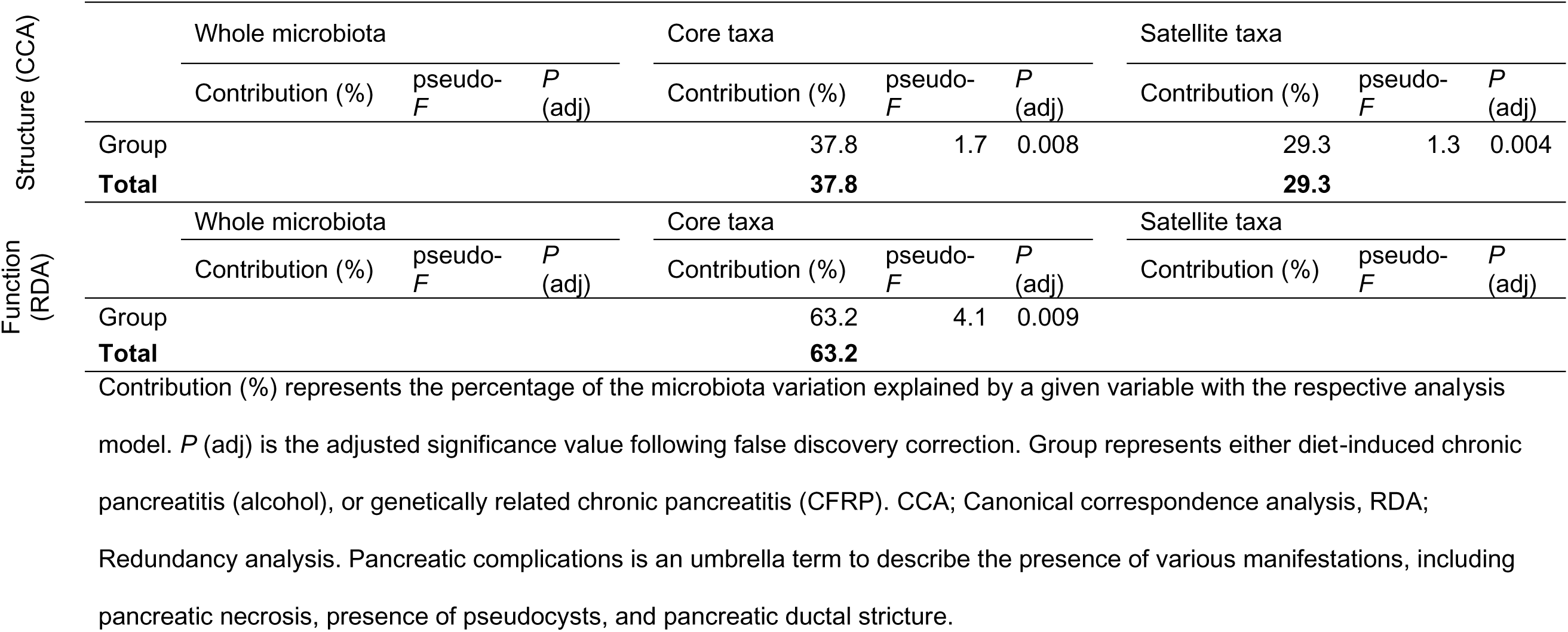
Ordination analyses to describe microbiota variation within the whole microbiota, core, and satellite taxa across chronic pancreatitis patients explained by significant clinical variables isolated from the forward stepwise-selection model.

**Table S8.**
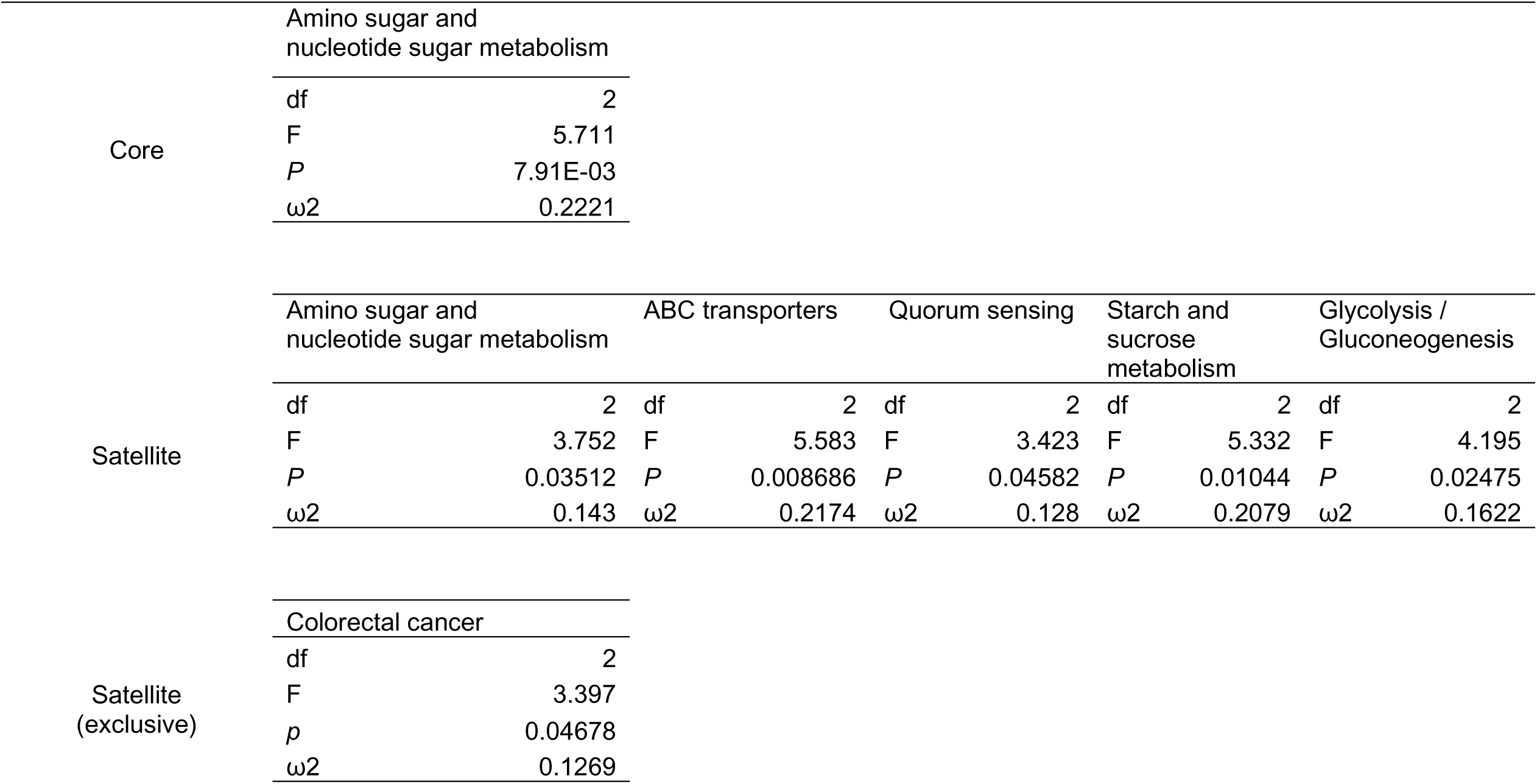
ANOVA summary statistics for KEGG level 3 hierarchical pathway comparisons across CLR-transformed relative abundance data.

**Fig. S2.**
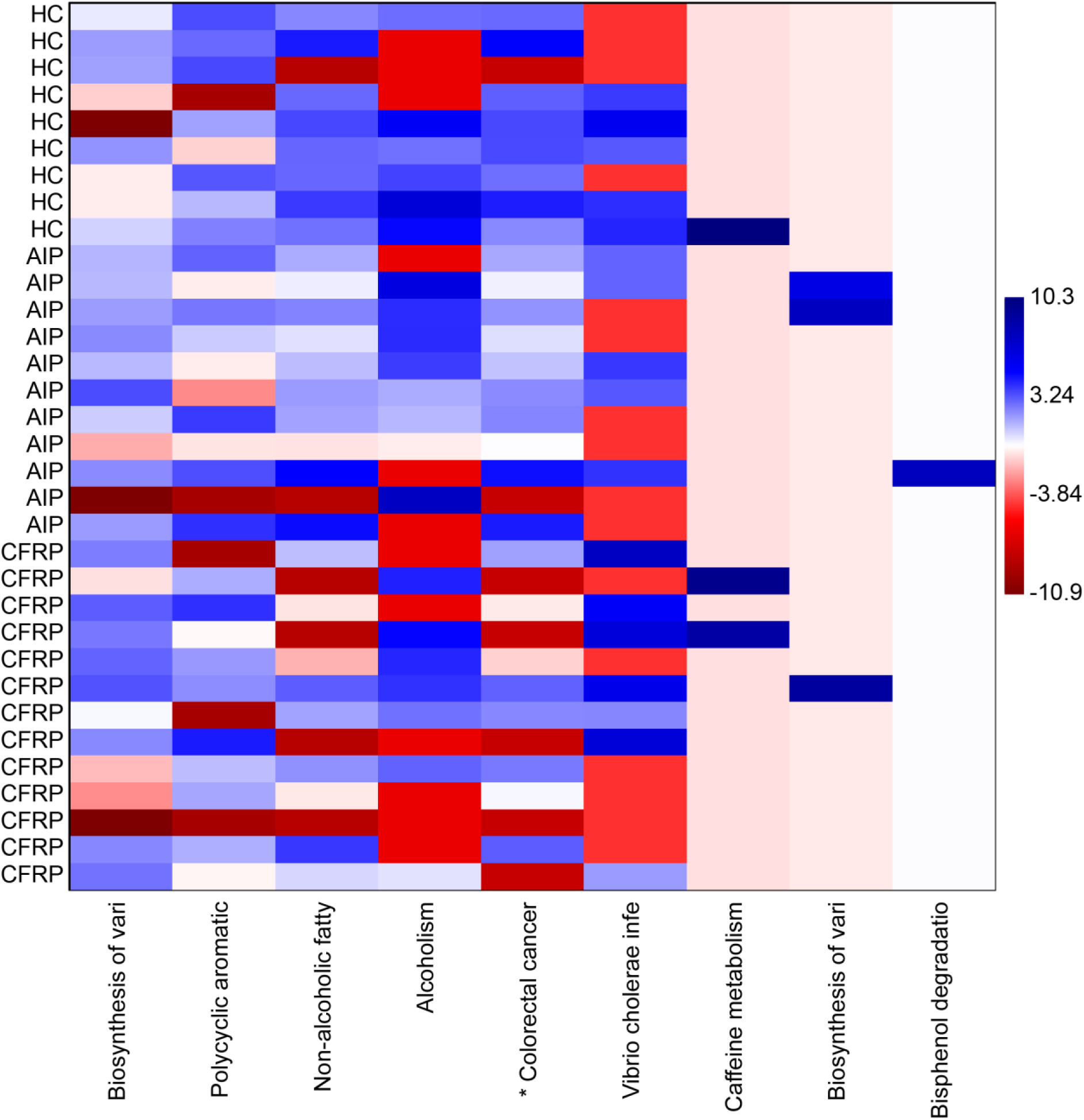
Differences in predicted metagenome function exclusive to the satellite taxa between study groups. CLR transformed compositional data was plotted to visualise differences among groups across KEGG level 3 hierarchical pathways relating to GI functionality and related physiological processes. Asterisks (*) denote significant differences across group means following ANOVA testing (*P* < 0.05). Full length pathway names and associated summary statistics can be found in Table S9.

**Table S9.**
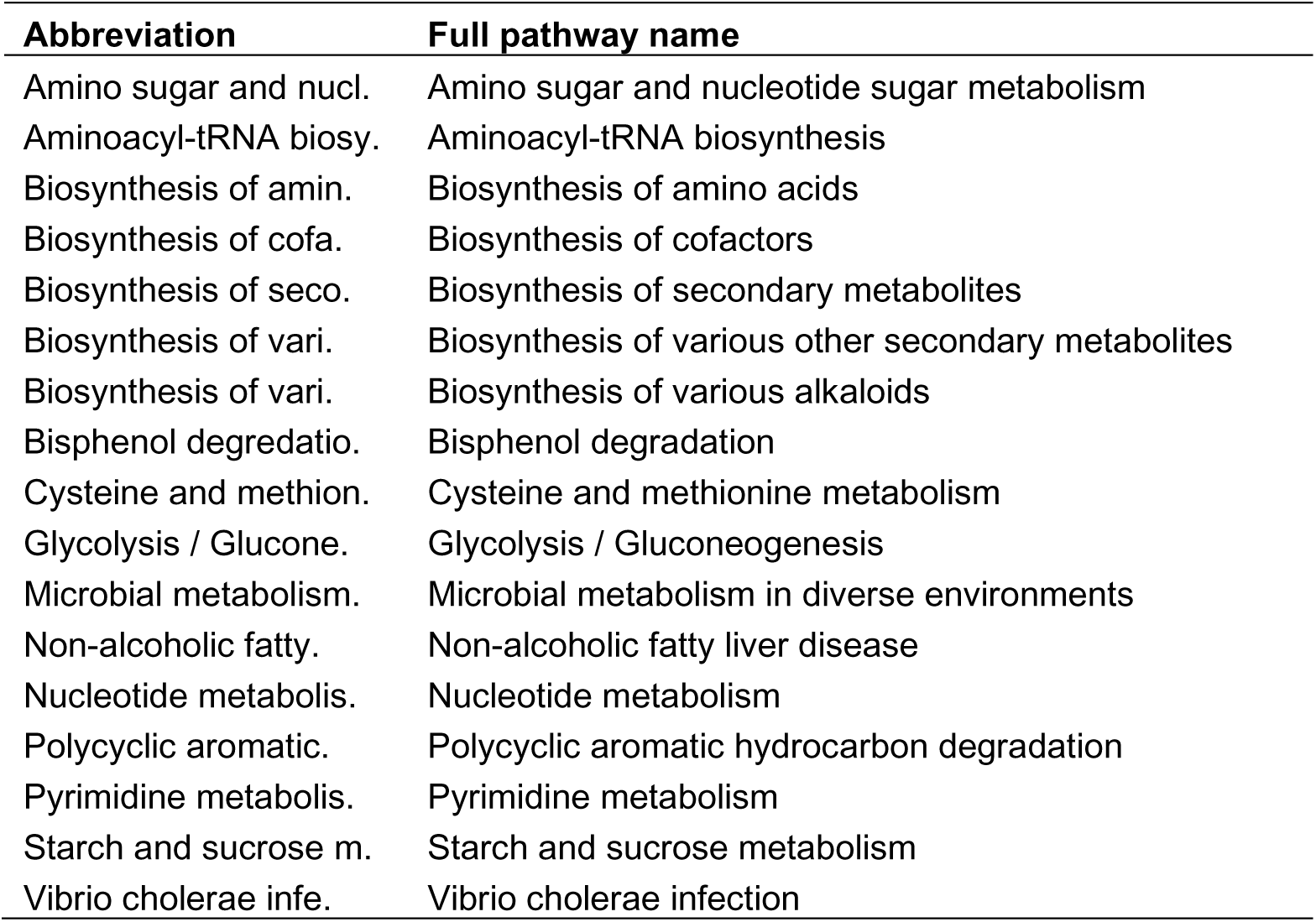
Details of KEGG level 3 hierarchical pathways.

